# Can computers understand words like humans do? Comparable semantic representation in neural and computer systems

**DOI:** 10.1101/843896

**Authors:** Linmin Zhang, Lingting Wang, Jinbiao Yang, Peng Qian, Xuefei Wang, Xipeng Qiu, Zheng Zhang, Xing Tian

## Abstract

Semantic representation has been studied independently in neuroscience and computer science. A deep understanding of human neural computations and the revolution to strong artificial intelligence appeal for a joint force in the language domain. We investigated comparable representational formats of lexical semantics between these two complex systems with fine temporal resolution neural recordings. We found semantic representations generated from computational models significantly correlated with EEG responses at an early stage of a typical semantic processing time window in a two-word semantic priming paradigm. Moreover, three representative computational models differentially predicted EEG responses along the dynamics of word processing. Our study provided a finer-grained understanding of the neural dynamics underlying semantic processing and developed an objective biomarker for assessing human-like computation in computational models. Our novel framework trailblazed a promising way to bridge across disciplines in the investigation of higher-order cognitive functions in human and artificial intelligence.

## 1 Introduction

Humans intuitively know that the meaning of the word *moon* is more related to *stars* than to *apples*. Establishing semantic similarity among concepts is a rudimentary adaptive trait for generalization. As an initial step for simulating human intelligence, computational models need to establish semantic relationship among words as well. To leap towards real artificial intelligence, we need to bridge representational formats independently developed from two complex systems – our brain and the computer.

Bridging the representational formats between computers and human brain has recently obtained promising breakthroughs. For example, in vision, the representations in visual hierarchy have been mapped onto distinct layers in deep neural networks (Khaligh-Razavi and Kriegeskorte 2014, Yamins et al. 2014). However, the important branch of artificial intelligence – natural language processing (NLP) – has yet to make substantial connections to higher-level cognitive function of language. The lack of fine-grained neurolinguistic processing models and granular neural recording methods constrains the progress in the language domain (Poeppel 2012). In this project, we proposed a novel approach to join forces across computer science and cognitive neuroscience. By searching for the correlations between neural activity recorded by electroencephalography (EEG) and semantic similarity learned by deep learning models of NLP, our work pioneered in bridging the gap in two ways. Specifically, (a) semantic information encoded in computational models unveiled the neural dynamics of semantic processing; (b) neural data quantified a temporal dynamic biomarker for objectively assessing human-like semantic similarity in NLP models.

### Semantics in computer science and cognitive neuroscience

Within computer science, semantic representation is the cornerstone of complex tasks such as information retrieval, question answering, machine translation, document clustering, etc. Earlier approaches were typically confined to algorithms that require the use of expert-knowledge-based corpus like WordNet (e.g., Resnik 1995, 1999, Lin 1998). Recent development in deep learning NLP models creates embedding representations based on the idea that lexical semantic information is reflected by word distribution (Harris 1954, Firth 1957, Miller 1986). Specifically, embedding models learn semantic representation from words’ distribution in their context in a large corpus. Distributional information of words is compressed into dense, lower-dimensional vectors. The similarity between two words can be represented by the cosine value of the angle between the vectors (see Turney and Pantel 2010, and see Fig. 1 for an illustration).

**Figure 1:**
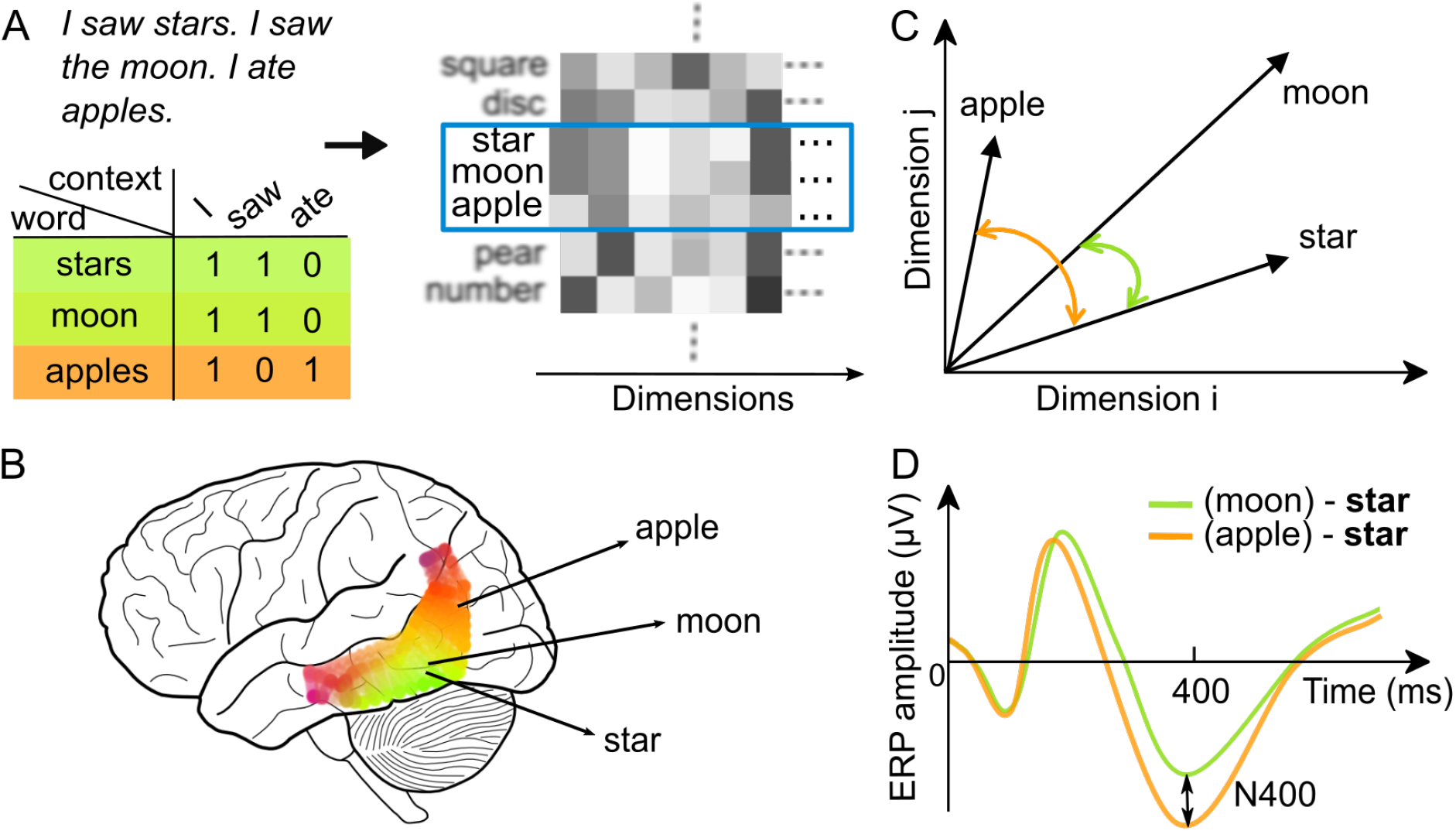
Schematic diagram of semantic representations in the human brain and word embedding models. **A)** A schematic diagram showing how the frequency of words in context yields embedding representations in computational models. Semantically similar words share higher distributional similarity, as illustrated by the counts of neighboring words in the sample mini corpus. Computational models learn semantic representation from words’ distribution and generate embedding representations. **B)** A schematic diagram showing the semantic space in the human brain à la Huth et al. (2016). Semantically more similar concepts are represented with more cortical overlaps, indicating shared features. **C)** A schematic diagram showing how the angle between high-dimensional vectors represents semantic similarity in computational models. The angle between two high-dimensional vectors (only two dimensions are used for demonstration) represents semantic similarity. The smaller an angle is (i.e., a higher cosine value), the higher semantic similarity (e.g., the angle between *star* and *moon* is smaller than the one between *star* and *apple*, because *star* and *moon* share more features, as shown in Fig. 1A). **D)** A schematic diagram showing how the N400 component of neural responses represent semantic similarity in the human brain (e.g., N400, see Kutas and Federmeier 2011). The less negative-going neural responses to a word are observed when it shares more semantic features with its preceding word (e.g., *star* shares more features with *moon* than with *apple*, as shown in Fig. 1B), yielding N400 effects.

Within cognitive science, ample empirical evidence has shown that the similarity of semantic representation has a profound impact on human behavior (Neely 1976, 1977, Lorch Jr 1982, Balota 1983, Anderson 1983, Roelofs 1992, Kiefer 2002). For example, in lexical decision, widely observed priming effects consist in that humans react to a word (e.g., *star*) faster when it is preceded by a semantically related word (e.g., in the pair *moon* – *star*) than by a semantically unrelated word (e.g., in the pair *apple* – *star*). The presentation of the first word (i.e., the prime) activates a node in a semantic network, which automatically spreads to neighboring nodes, facilitating the processing of the second word (i.e., the target) if it is semantically related (Collins and Loftus 1975)

These behavioral findings on semantic similarity were further supported by neuroimaging studies. For example, the voxel-wise modelling neuroimaging study has yielded a semantic map in the human brain, on which concepts sharing more semantic features are mapped to closer brain areas (Huth et al. 2016). In electrophysiological studies, N400 effects – less neural activity around 400 ms after the onset of a more semantically expected word – were observed in both contextual and priming settings (e.g., Bentin et al. 1985, Kutas and Hillyard 1989, Holcomb 1993, Brown and Hagoort 1993, Federmeier and Kutas 1999, Deacon et al. 2000, Kiefer 2002) (see Fig. 1).

So far semantic representations have been investigated independently in computer science and cognitive neuroscience. It remains unclear to what extent representations yielded from computer models resemble to the ground truth of human representation.

### Bridging semantic representations in the human brain and computer models

Recently, computational models have started advancing our understanding of language processing in human brain(Brennan 2016). The bridging of representations between neural activity and computational models has been preliminarily investigated in sentential context using N400 effects (Ettinger et al. 2016, Broderick et al. 2018). However, neural activity recorded during the comprehension of sentential stimuli and continuous speech was driven by both compositional processing (e.g., the composition between *lamb* and *stew*, yielding *lamb stew*, see e.g., Bemis and Pylkkänen 2011, Zhang and Pylkkänen 2015, Pylkkänen 2019) and semantic processing (e.g., similarity-based spreading from *lamb* to *stew*), making neural data hardly comparable with pure semantic representations yielded from word embedding models.

Therefore, our study focused on the representation of lexical semantics in the human brain and computer models. We adopted a canonical semantic priming design that elicited the measures of semantic similarity in the brain, directly comparable to semantic representations yielded by computational models without confounding factors from compositional processing. We predicted that the two measures from the brain and computers would correlate in a rather narrow time window within classical N400 component, presumably at the beginning of the processing purely related to semantic representation without contamination from compositional processing.

Moreover, we selected three representative word embedding models, differing in the way of learning semantic representation. The CBOW (Continuous Bag-of-words) model (Mikolov et al. 2013) solely uses local context – a number of words immediately preceding and following a word. The other two models are based on CBOW. The GloVe (Global Vectors) model (Pennington et al. 2014) combines both local context and global corpus statistics for learning word representation. The CWE (Character-enhanced Word Embedding) model (Chen et al. 2015) captures both word-external local contextual information and word-internal character information. We predicted that both GloVe and CWE would correlate with brain responses better than CBOW. The better correlation would occur at different times because of particular features of the models – CWE at an earlier perceptual stage due to its inclusion of character-level information, whereas GloVe at a later stage reflecting semantic processing.

By assessing the representational formats with a well-controlled experiment and millisecond-level neural recordings, we provided a framework directly bridging semantic representations between the human brain and computers. Our aim was twofold: (a) information encoded in NLP models contributed to a finer-grained understanding of the neural dynamics underlying semantic processing, especially pinning down specific time windows for known components involved in the semantic processing by the brain; (b) neural data contributed an objective, temporal dynamic, assessment for human-like language processing in NLP models.

## 2 Methods

### 2.1 Participants

A group of 30 healthy right-handed native Chinese speakers participated in the study. Participants were recruited with the use of online flyers. All had normal or corrected-to-normal vision. The EEG recordings of the 30 participants took place from March 12th, 2018, to October 7th, 2018. Five participants were excluded from data analyses: three due to excessive noise during recording, and two for being outliers in terms of accuracy in the behavioral task (more than 3 standard deviations below the average). Thus, 25 participants were included in EEG data analyses (14 females; average age = 22.6 years, *SD* = 2.8 years). All data were collected at the EEG lab at the NYU-ECNU Institute of Brain and Cognitive Science at NYU Shanghai (Shanghai, China). This study was approved by the local ethical committee at NYU Shanghai. Written consents were obtained from each participant. No participants declined participation during or after data collection, and given the experimental design, participants were not allocated into groups.

### 2.2 Experimental design and stimuli

Our EEG experiment adopted a canonical two-word priming paradigm, with stimuli visually presented to the participants. We used 240 pairs of two-character Chinese nouns as critical stimuli. We randomly selected nouns to form ‘prime-target’ pairs. Among these ‘prime-target’ word pairs, some pairs (e.g., 月亮 (moon) – 星 星 (star)) are intuitively of a higher semantic similarity than others (e.g., 苹果 (apple) – 月亮 (moon)). This random selection procedure yielded a distribution of semantic similarity (between prime and target) shown in Fig. 2B (see the entire stimuli list at https://ray306.github.io/brain_NLP/).

**Figure 2:**
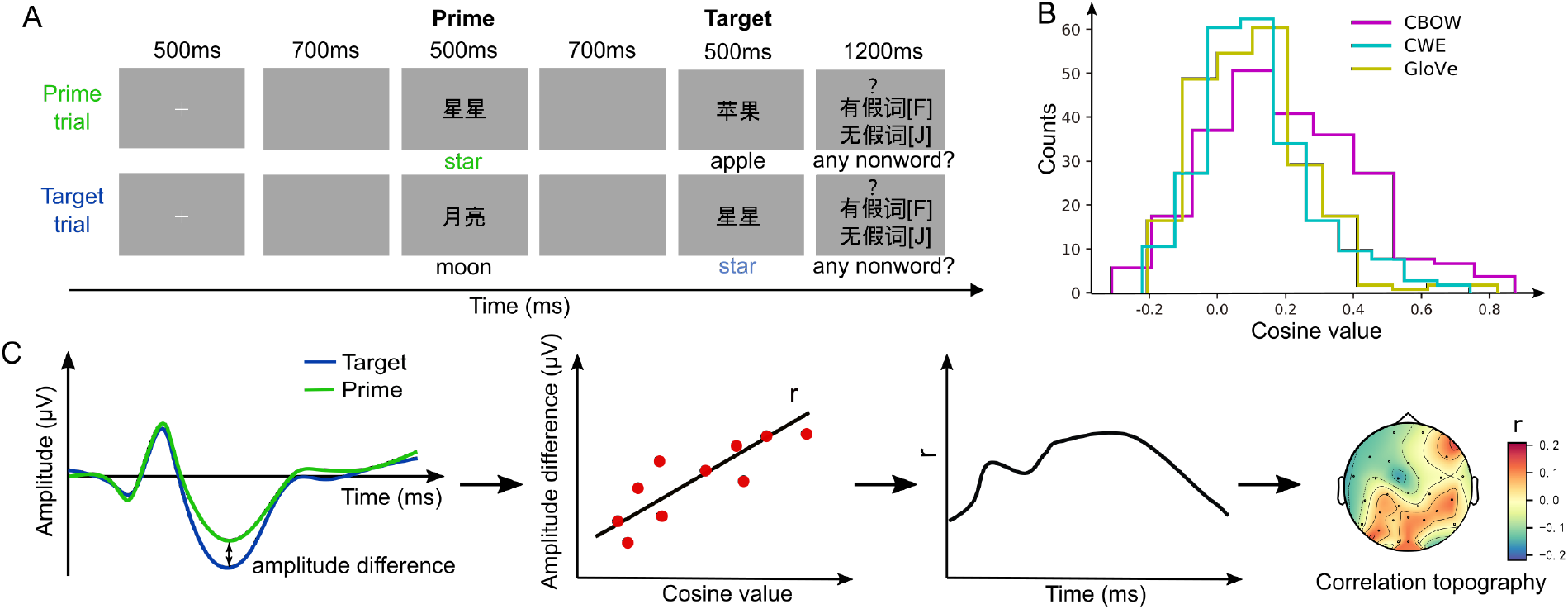
Experimental procedure and single-trial correlation analysis. **A)** The trial structure of the experiment. Sample trials are illustrated for the two-word priming paradigm. In each trial, a prime word was followed by a target word. Each word (here 星星(star)) was used once at the prime position (in the prime trial) and once at the target position (it the target trial). English translations below the screens are for demonstration only, but not included in the expreiment. **B)** Stimuli statistics of semantic similarity generated by the three computational models. **C)** The flowchart of single-trial correlational analysis, (i) Computing the amplitude differences between single-trial EEG responses to the same word at its target vs. prime presentation (target minus prime); (ii) For the 240 word pairs, calculating the Pearson’s correlation between cosine values generated from computational models and EEG response differences from step (i) at each time point in each sensor; (iii) The obtained Pearson’s correlation coefficients form a waveform across time for each sensor; (iv) The distribution of Pearsons’s correlation coefficients from all sensors is plotted in a topography at each latency.

To construct 240 critical trials, we used 240 distinct nouns. Each noun appeared at the prime position once and at the target position once (see Fig. 2A). For each noun (e.g., 月亮 (moon)), the EEG responses elicited at the prime position (e.g., in the trial 月亮 (moon) – 星星 (star)) represent semantic retrieval of its out-of-context meaning. Whereas, the EEG responses elicited at the target position (e.g., in the trial 苹果 (apple) – 月亮 (moon)) include the influence of the preceding word. Thus, the difference between these two EEG responses to the same word at different positions is priming effects, reflecting semantic similarity without the contamination from semantic retrieval. Therefore, we extracted the neural measure directly comparable to the semantic similarity computed from NLP models. Moreover, we extended the previous condition-level computation of ERP differences to the trial-level and provided a trial-level measurement of semantic priming effects.

We added 120 additional pairs of stimuli as fillers, in which either the prime or the target was a two-character non-word (e.g., 害天, 粽七). Thus, a total of 360 trials were included in this experiment. Participants were instructed to perform a lexical decision task, judging whether a trial contained a non-word. The purpose was to keep participants alert, encouraging them to process the stimuli at least to the lexical semantics level.

The trial structure is illustrated in Fig. 2A. Each trial started with a fixation lasting for 500 ms. After a 700 ms blank screen, the prime was presented for 500 ms. After another 700 ms blank screen, the target was also presented for 500 ms, followed by a question mark ‘?’ and a prompt for the lexical decision task. The stimuli were in a white 40-point Songti font on a gray background. The 360 trials were divided into 6 blocks, each containing 60 trials. The critical trials and fillers were pseudo-randomized and quasi-evenly distributed in each block. The blocks were also pseudo-randomized. Between blocks, participants could take a short rest. The experimental presentation was programmed with a Python package – Expy (https://github.com/ray306/expy), an in-house software for presenting and controlling psychological experiments, available at http://slang.science.

### 2.3 Procedure of data collection

in an electrically-shielded and sound-proof room. EEG data were continuously recorded via a 32-channel ActiChamp system (Brain Products). Electrodes were held in place on the scalp by an elastic cap (ActiCap) in a 10-20 configuration as shown in Fig. 3A. Two more electrodes were placed below the left eye and at the outer canthus of the right eye to monitor vertical and horizontal eye movements (electro-oculogram, EOG). Impedance was kept less than 10 kΩ for all electrodes. The EEG signal was recorded in single DC mode, digitized at a sampling rate of 1000 Hz and online referenced to the vertex (Cz), with the use of the software BrainVision PyCoder. The recording session lasted approximately 30 minutes.

**Figure 3:**
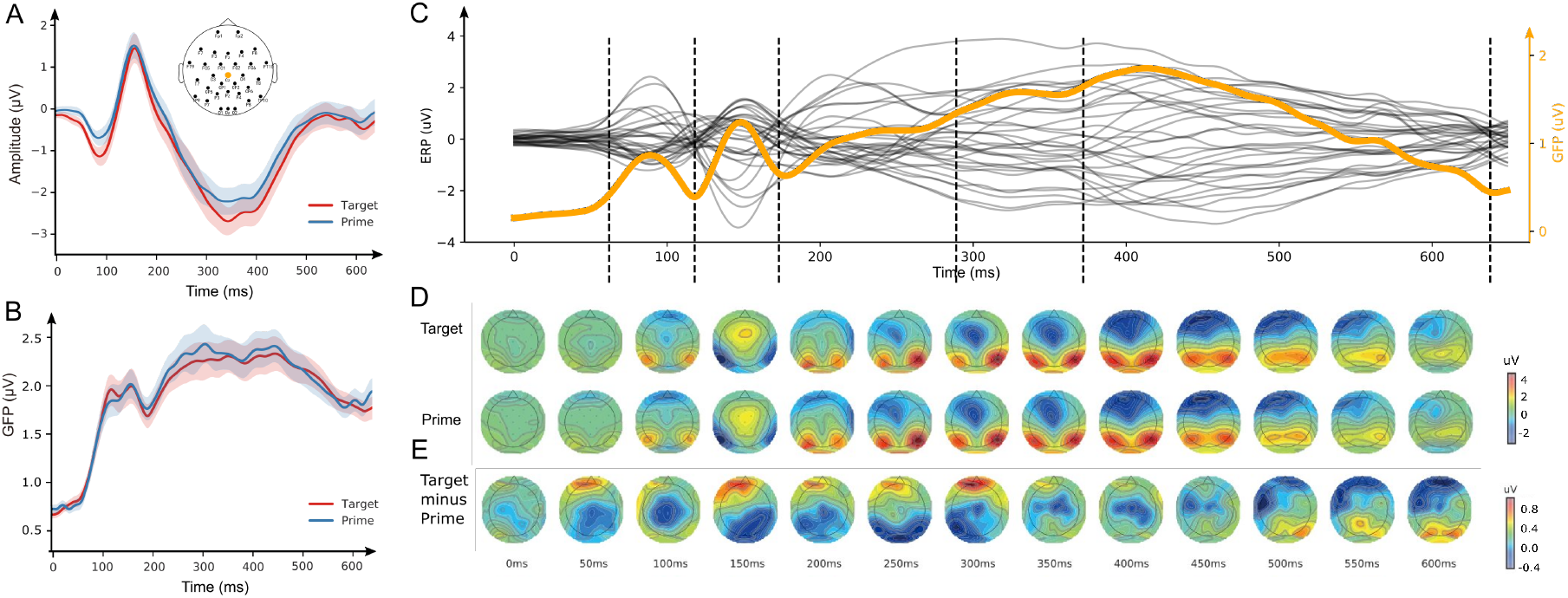
Event-related waveform and topographic responses consistent with perceptual and semantic processes in language comprehension. **A)** The waveform responses in a representative channel (Cz). Typical N400 profile was observed for processing a word at both prime and target positions. The montage of sensor locations is inserted with the selected channel Cz highlighted. **B)** The dynamics of GFP. The aggregated neural activity across all sensors represented in GFP shows the similar dynamics that has clear perceptual and semantic activation. **C)** The temporal components revealed in the grand averaged ERP responses across targets and primes. Each black line represents ERP responses in each channel. The orange line represents the GFP across all sensors. The vertical dashed lines label the temporal boundaries between ERP components revealed by an automatic segregation method. **D)** The temporal progression of topographies. The topographies for target and prime were represented in the upper and lower rows respectively. Similar topographic patterns and temporal progressions were observed for processing a word at both target and prime positions. **E)** The temporal progression of topographic differences. Differences resulted from subtracting prime from target revealed classic N400 topographic patterns from 250 to 600 ms.

### 2.4 Data pre-processing

Only the 240 critical trials were included in EEG analysis. EEG data were processed and analyzed with EasyEEG toolbox version 0.8.4.1 (Yang et al. 2018, https://github.com/ray306/EasyEEG). Raw EEG data were bandpass filtered between 0.1 and 30 Hz and epoched from 200 ms before to 800 ms after the onset of a word. Epochs were baseline corrected with the 200 ms interval before word onset. We removed those epochs affected by large vertical or horizontal eye movements, based on data recorded from the two electrodes monitoring EOG. We further visually inspected the epochs and removed those with large artifacts. The data were re-referenced to the average reference.

### 2.5 Data analyses

#### 2.5.1 Behavioral data

We checked the accuracy and reaction times for all 360 trials. Reaction times were measured from the onset of prompt for each trial and for each participant. We ran a two-tailed *t*-test on the data of accuracy and reaction times between critical trials and fillers, to verify whether participants paid attention to the stimuli.

#### 2.5.2 EEG data

The analysis of EEG data constituted two parts. The first part aimed to examine the validity of the data by checking the ERP components in reading as well as N400 priming effects with the use of data averaged across trials (see Section 2.5.2.1). The second part was at the trial level, aiming to test (a) whether EEG responses can be predicted by a computational model within the typical time window for N400 priming effects (see Section 2.5.2.2) and (b) among CBOW, GloVe, and CWE, which computational model was the best predictor at which time point (see Section 2.5.2.3).

##### 2.5.2.1 ERP analysis

Trials were averaged for prime and target respectively. We plotted the ERP waveforms in a representative channel (Cz) for ERP to compare our data with N400 effects reported in literature. To summarize and visualize the distributed energy fluctuation, we plotted the dynamics of Global Field Power (GFP, see Lehmann and Skrandies 1980), calculated as a geometric mean of electric potentials across all sensors. To reveal and visualize ERP components during word processing, we used an automatic segregation method (Topography-based Temporal-analysis Toolbox, TTT) to detect component boundaries and plotted boundaries along with average ERP responses of each channel and GFP (Wang et al. 2019). To visualize the dynamics of activation patterns, we plotted the topographies across time for ERP responses to prime and target as well as the differences between the two (i.e., target minus prime).

##### 2.5.2.2 EEG data analysis at trial level (a): testing whether EEG responses can be predicted by a computational model

All the three selected word embedding models (i.e., CBOW, GloVe, and CWE) were trained on Chinese Wikipedia. These models calculated cosine similarities for the 240 word pairs used as critical stimuli, and we correlated the model-generated cosine similarities with single-trial EEG responses, according to the following procedure (see Fig. 2C):

First, for each word, we subtracted the EEG responses to its presentation at the prime position from those responses at the target position. This EEG difference for each word represents priming effects with no contamination of semantic retrieval.

Second, we calculated the correlation co-efficient *r* between ERP differences (computed from 240 critical trials by Step 1) and model-generated semantic similarities (cosine values). This calculation of correlation was performed at each time point in each channel.

Third, the correlations of all time points at a channel yielded a temporal progression of correlations at this channel.

Fourth, based on the previous three steps, we calculated the temporal progression of correlations for all channels and obtained a series of topographies of correlations along the time course.

We obtained a null distribution of *r* values by shuffling the pairing among the 240 EEG response differences and the 240 cosine similarities for 1000 times. Empirical *r* values were checked against this null distribution to determine the statistic significance (at the level of *p <* 0.05) at each time point.

##### 2.5.2.3 EEG data analysis at trial level (b): testing which computational model was the best predictor at which time point

When testing which word embedding model (among CBOW, GloVe, and CWE) was the best predictor at which time point, we conducted permutation tests on correlation *r* values averaged across channels to estimate the overall predictability of each model. We did the same permutation tests on correlation *r* values for each channel to examine the spatial distribution of the predictability of each model.

Specifically, from the correlation between EEG responses in each of the 32 channels at each of the 800 milliseconds and cosine similarities computed from each of the three computational models, we obtained a 32 *×* 800-dimensional matrix of *r* values for each model.

To estimate the overall predictability of each model, we averaged the absolute *r* values across channels, yielding a line of temporal progression of *r* for each word embedding model. At each time point, we randomly shuffled the pairing between EEG responses and cosine values generated by the three models for 1000 times. The shuffling yielded a null distribution of *r* differences between any two models. Empirical *r* differences were checked against this null distribution at each time point.

We did the same permutation tests for each channel to further compare the predictability of models and investigate the site of effects.

## 3 Results

### 3.1 Behavioral data

The mean accuracy of lexical decision task was 94.6% (SD = 2.4%). The two-tailed *t*-test revealed significant differences between critical trials and fillers (mean accuracy and SD for critical trials: 96% (2.5%); mean accuracy and SD for fillers: 91% (4.6%); *t* (24) = 4.99; *p <* 0.001).

The mean reaction time was 289 ms (SD = 102 ms). The two-tailed *t*-test also revealed significant differences between critical trials and fillers (mean reaction time and SD for critical trials: 296 ms (103 ms); mean reaction time and SD for fillers 274 ms (101 ms); *t* (24) = 4.475; *p <* 0.001).

Behavioral data indicated that participants reacted differently towards critical trials and fillers, suggesting that they fully processed lexical semantic information.

### 3.2 EEG data

#### 3.2.1 Results from ERP analysis

ERP responses were obtained after averaging trials for prime and target respectively (Fig. 3). The waveform ERP responses at a representative channel, Cz, clearly indicate the evolution of ERP components associated with reading a word (Fig. 3A). Responses to both target and prime showed early visual responses N1 and P2 as well as semantics-related N400 component, consistent with well-established literature (Kutas and Federmeier 2011). Similar evolution of ERP components was also observed in the dynamics of GFP which included activity of all sensors (Fig. 3B), demonstrating the reliability of elicited data without the potential pitfalls of subjective bias. The boundaries of ERP components were detected based on an automatic segregation method (Wang et al. 2019) and plotted in Fig. 3C. The component after visual processing was further segregated into three sub-components.

Topographic responses to prime and target demonstrate consistent evolution of response patterns (Fig. 3D), suggesting common cognitive functions unfolding over time during the reading of these words at prime and target positions. Topographic differences between target and prime showed magnitude differences in sensors over frontal and temporo-parietal regions around 300 ms (Fig. 3E), consistent with the pattern of typical N400 priming effects (see Kutas and Federmeier 2011)

Our ERP responses were temporally and spatially consistent with well-established N400 priming effects, demonstrating the reliability and validity of neural measures on semantic similarity.

#### 3.2.2 Results from trial-level analysis (a): single-trial EEG responses can be predicted by a computational model

We selected GloVe as a representative NLP model. The generated measure of semantic similarity was correlated with single-trial EEG response differences between prime and target (Fig. 4). The correlation was significant at 300 ms after word onset at channel Oz: *r* = 0.173 (*p* = 0.007) (Fig. 4A). The dynamics of *r* was obtained in the same channel (Fig. 4B). A non-parametric statistics revealed that the GloVe-generated semantic similarity values significantly correlated with EEG response differences between 226 and 306 ms.

**Figure 4:**
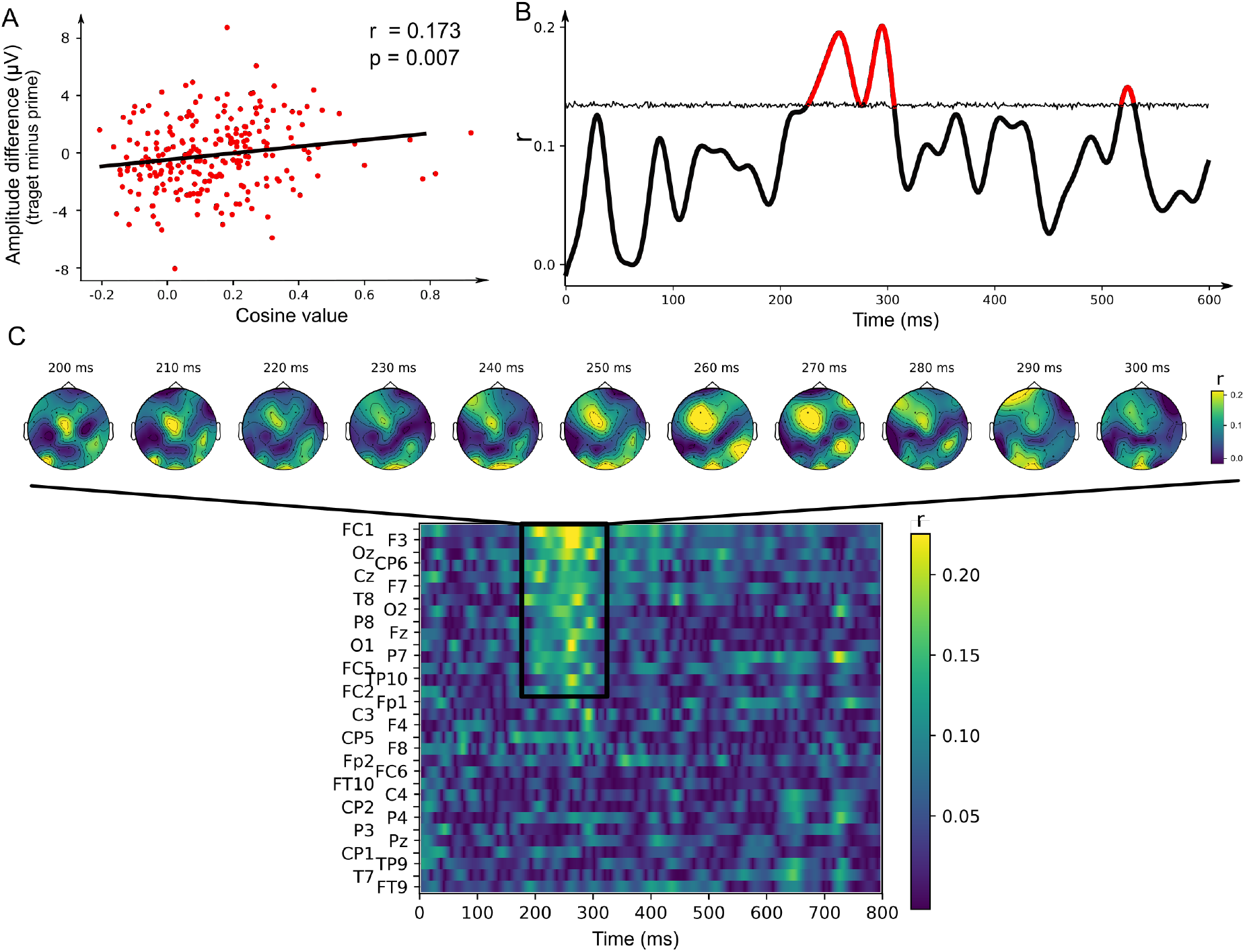
Correlations between EEG responses and a word embedding models reveals the dynamics of semantic processing. **A)** Significant correlation was observed between EEG responses in channel Oz at the latency of 300 ms and cosine values computed by the model GloVe. **B)** The temporal progression of correlations (channel Oz). Significant correlations were observed between 226 and 274 ms, between 279 and 306 ms, and between 518 and 529 ms (in red). The significance was determined by the threshold (horizontal line) obtained in a non-parametric permutation test at each time point (alpha level at 0.05). **C)** The spatio-temporal characteristics of correlations. The heatmap of correlations across time and channels revealed significance between 200 and 300 ms in about half of the sensors. The progression of topographies in the time window of significance is zoomed in above. Significant correlations were concentrated in the sensors above the left frontal and tempo-parietal regions.

The spatial distribution of *r* value was further investigated, by computing the correlations in all sensors (Fig. 4C). The heamap shows that correlations in about half of the sensors were significant between 200 to 300 ms, consistent with the results in Fig. 4B. The distribution of significant correlations in this time window was scrutinized by delineating the evolution of topographies. Most robust correlations were found at sensors over the left frontal and occipital regions, consistent with the typical pattern of N400 effects. The observed semantic processing in a narrow and early time window was consistent with the findings of semantic dynamics in ERP responses after removing temporal variance among trials (Wang et al. 2019). Taken together, these results demonstrated that NLP models can predict EEG responses, suggesting the common semantic representations between two complex systems.

#### 3.2.3 Results from trial-level analysis (b): The three NLP models distinctively correlated with EEG responses

We compared the predictability of three selected NLP models (CBOW, GloVe, and CWE) with permutation tests along the temporal progression. Averaged *r* values across channels in any two of the three models were subject to pairwise comparisons. The results revealed three time windows (lasting for at least 10 ms) within which one model was a significantly better predictor than another one at each time point: (a) CWE predicted significantly better than GloVe between 94 and 122 ms; (b) GloVe predicted significantly better than CWE between 244 and 256 ms; (c) GloVe predicted significantly better than CBOW between 202 and 251 ms. GloVe was a significantly better predictor than the other two models between 244 and 251 ms (yellow shaded area in Fig. 5A).

**Figure 5:**
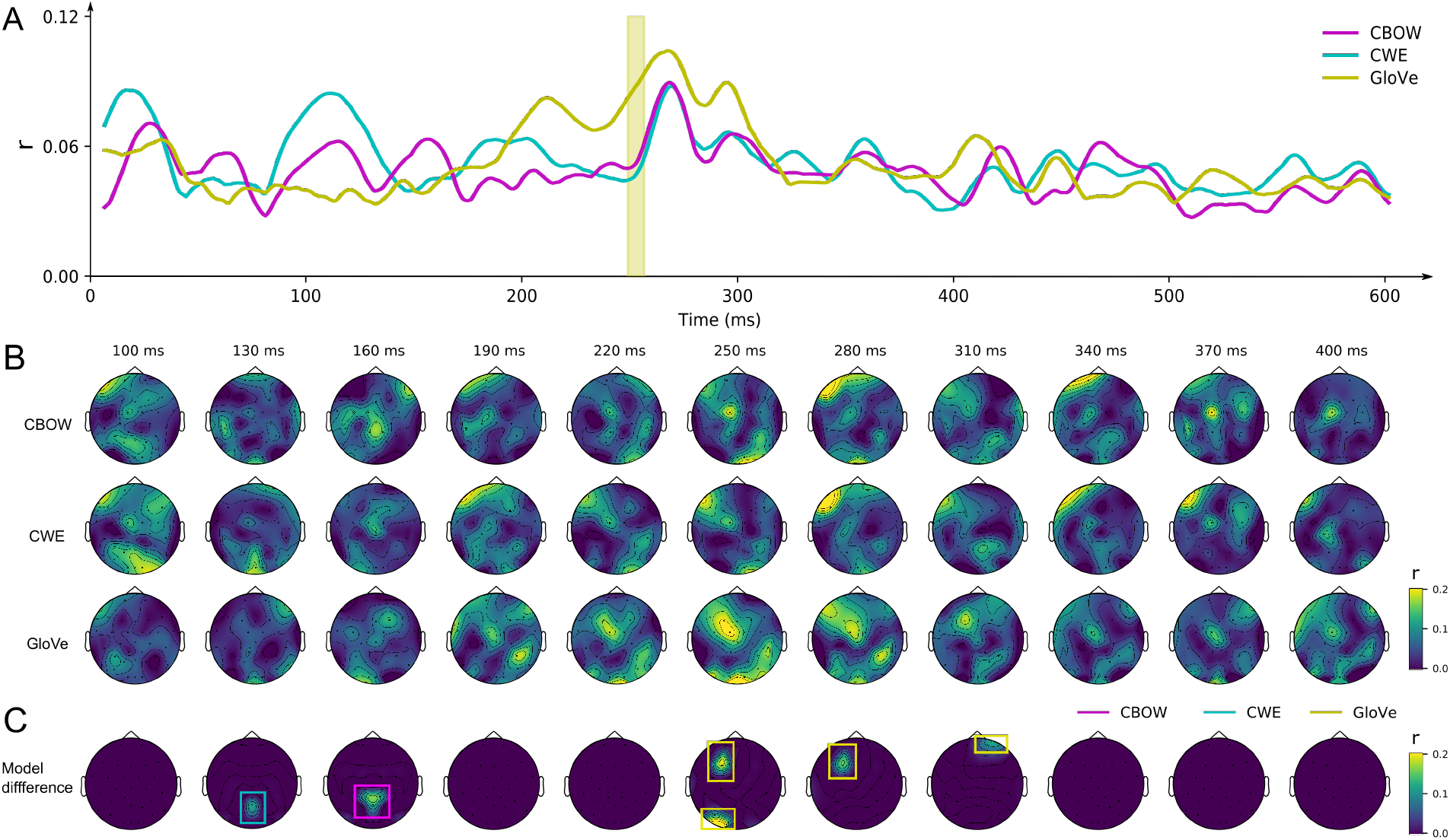
Three word embedding models distinctively correlate with EEG responses. **A)** The temporal progression of averaged correlations across sensors for each computational model. The correlation for GloVe was significantly better than the other two models between 244 and 251 ms, as highlighted in the shaded window. The significance was determined by non-parametric permutation tests. **B)** The temporal progression of correlation topographies for each computational model. Similar patterns were observed among all models. **C)** The tempo-spatial characteristics of correlation differences among the three computational models. Pairwise non-parametric permutation tests in each sensor revealed distinct predictability at different latencies for each model.

The topographies of *r* values for all three models were plotted in Fig. 5B, demonstrating that the correlation patterns were spatially consistent among the three models. In particular, high correlations were observed in sensors over the left frontal and occipital regions around 250 to 300 ms, similar as the observation in Fig. 4. The similar spatio-temporal configuration was obtained in permutation tests at channel level, which further revealed that GloVe was the best predictor at sensors over the left frontal and occipital regions around 250 to 300 ms (Fig. 5C). These consistent results in temporal and spatial domains provide strong evidence for the dynamics of semantic processing.

Moreover, CWE was the best predictor around 130 ms in posterior channels. Consistent spatio-temporal configurations for this earlier effect were also observed across all the three models (Fig. 5B). CBOW was the best predictor around 160 ms in posterior channels. Taken together, these results show that the three word embedding models distinctively correlated with ERP differences at distinct latencies.

## 4 Discussion

In this study, we investigated whether and how the lexical semantic representation that independently established in the human brain and computational models share similar formats. We found that semantic similarities computed by word embedding models correlated with EEG semantic priming responses in an early and narrow time window of N400 component. Moreover, distinct word embedding models that include different weighting of orthographic and semantic information correlated with neural responses at perceptual and semantic processing stages. Our study provided strong evidence suggesting that the dynamic processing of lexical semantics can be characterized by word embedding models based on the commonality of semantic representation between two complex systems.

With a better controlled two-word semantic priming paradigm and non-invasive electrophysiological recordings, we provided an analytical approach to collaboratively investigate the semantic representations in two independent complex systems. Computational models can yield quantitative hypothesis to investigate neural processing, and neuroscience data can back-feed to computer models towards creating a stronger artificial intelligence that better emulate neural processes and human behavior. The current study provided a novel framework on how cognitive neuroscience and computer science can be bridged in a bi-directional investigation of the computational mechanisms in language research.

Computer science can help investigating neuroscientific theories. Granular aspects of linguistic information, such as lexical semantics, can be captured by computational models precisely, without contamination from other factors. Such dedicated and quantitative linking hypothesis between computers and brain provides lens to scrutinize neural computations. The millisecond-by-millisecond single-trial correlational analysis in the current study strikingly narrowed down the time window associated with well-established N400 component that commonly lasts from 250 to 600 ms after a word onset. The observation of significant correlation in a narrow and early time window remarkably reflected the processing of lexical semantics per se. These results can resolve a long lasting debate regarding to one of the most investigated linguistic processing components, N400 – whether it is integration (e.g., Hagoort et al. 2004) or semantic retrieval (Kutas and Federmeier 2011). Our results based on semantic representation extracted independently from computational models suggest that the commonly observed long duration of N400 presumably contains several sub-processes, and semantics-related processing starts at the beginning.

Neuroscience can facilitate the journey to strong artificial intelligence. The current study advances in this direction from three aspects. First, neural measures can provide a biomarker for objectively assessing android performance of computational models. The correlations between two complex systems vary as a function of model selections (Fig. 5A). The model GloVe correlated with neural data significantly better at around 250 ms than the other two models, suggesting that the implementation of global context yielded more human-like semantic representation. Second, the characteristics of neural dynamics can dissect computational models to probe their features. Distinct models showed better correlations at different latencies (Fig. 5C), suggesting CWE that correlated best at around 130 ms weighted more on lexical-orthographic features, whereas GloVe weighted more on lexical semantics.

Third, this study trailblazes a database that will integrate research communities that vary across disciplines, cultures, and societies (https://ray306.github.io/brain_NLP/). The database can help computer scientists to evaluate how human-like their models are and to assess in which aspects the human-like features are. Moreover, the obtained millisecond-level, continuous neural data can help improve model performance and generalize across tasks by optimally integrating the best aspects of models based on dynamic featural processing. Our database (currently only in Mandarin Chinese and English) is expected to expand to many other languages and dialects. We welcome the whole research community to contribute. This joint force will broaden the horizon and provide a unique opportunity to generalize computational models for language processing.

Relating AI models and cognitive neuroscience has brought fruitful findings in other domains of cognition. For example, in vision, the state-of-the-art works by Kriegeskorte’s and DiCarlo’s groups (Kriegeskorte and Kievit 2013, Khaligh-Razavi and Kriegeskorte 2014, Yamins et al. 2014) have established a mapping between features in different layers of deep neural network model and neural representation in the hierarchical processing in the brain. Our current study was an attempt to create such mapping in the domain of language. Unlike research in vision that can obtain from animal models using invasive methods, linking NLP models and language processing in human brain is constrained by the limits of neuroimaging methods. We carefully chose semantic features and a functional paradigm that can establish direct mapping between computational models and human brain in the linguistic domain. This endeavor opened a brand new door towards a full understanding of computational mechanisms of language processing in both complex systems.

## 5 Conclusion

By investigating the representational formats of comparable lexical semantic features between complex systems with fine temporal resolution neural recordings, we provided a novel framework directly bridging neuroscience and computer science in the domain of language. This framework brought a finer-grained understanding of the neural dynamics underlying semantic processing and developed an objective biomarker for assessing human-like computation in NLP models. Our study suggested a promising way to join forces across disciplines in the investigation of higher-order cognitive functions in human and artificial intelligence.

## Acknowledgements

We thank Yuqi Hang, Yunyun Shen, Jiaqiu Sun, and Hao Zhu for their kind help during data collection. This research was funded by National Natural Science Foundation of China (Grant No. 31871131 to X.T.), the Science and Technology Commission of Shanghai Municipality (Grant No. 17JC1404104 to X.T., Grant No. 17JC1404101 to Z.Z., and Grant No. 17JC1404103 to X.Q.), as well as the Program for Eastern Young Scholar at Shanghai Institutions of Higher Learning (to L.Z.).

## Ethical statement

This study was approved by the local ethical committee at NYU Shanghai. Written consents were obtained from each participant.

## Data availability

(i) Word pairs used in EEG data collection, (ii) raw and processed EEG data, and (iii) cosine similarities computed from the three word embedding models (CBOW, CWE, and GloVe) can be downloaded from https://osf.io/tdrfw/. These data and their detailed description can also be found at https://ray306.github.io/brain_NLP/.

## Code availability

The open source EEG data acquisition software of BrainVision PyCoder can be found at https://www.brainproducts.com/downloads.php?kid=38. The Python package used for stimuli presentation is available at https://github.com/ray306/expy.

The open source code of the model CBOW is available at https://github.com/rare-technologies/gensim.

The open source code of the model CWE is available at https://github.com/Leonard-Xu/CWE.

The open source code of the model GloVe is available at https://github.com/stanfordnlp/GloVe.

The EEG data preprocessing code is based on the EasyEEG toolbox version 0.8.4.1 (Yang et al. 2018, https://github.com/ray306/EasyEEG). Specific codes used for EEG data (pre)processing, cosine similarity computation, and correlation analyses can be downloaded from https://osf.io/tdrfw/.

## References

Anderson, John R. 1983. A spreading activation theory of memory. Journal of verbal learning and verbal behavior 22:261–295.

Balota, David A. 1983. Automatic semantic activation and episodic memory encoding. Journal of verbal learning and verbal behavior 22:88–104.

Bemis, Douglas K, and Liina Pylkkänen. 2011. Simple composition: A magnetoencephalography investigation into the comprehension of minimal linguistic phrases. Journal of Neuroscience 31:2801–2814.

Bentin, Shlomo, Gregory McCarthy, and Charles C. Wood. 1985. Event-related potentials, lexical decision and semantic priming. Electroencephalography and clinical Neurophysiology 60:343–355.

Brennan, Jonathan. 2016. Naturalistic sentence comprehension in the brain. Language and Linguistics Compass 10:299–313.

Broderick, Michael P, Andrew J Anderson, Giovanni M Di Liberto, Michael J Crosse, and Edmund C Lalor. 2018. Electrophysiological correlates of semantic dissimilarity reflect the comprehension of natural, narrative speech. Current Biology 28:803–809.

Brown, Colin, and Peter Hagoort. 1993. The processing nature of the N400: Evidence from masked priming. Journal of cognitive neuroscience 5:34–44.

Chen, Xinxiong, Lei Xu, Zhiyuan Liu, Maosong Sun, and Huanbo Luan. 2015. Joint learning of character and word embeddings. In Twenty-Fourth International Joint Conference on Artificial Intelligence.

Collins, Allan M., and Elizabeth F. Loftus. 1975. A spreading-activation theory of semantic processing. Psychological review 82:407.

Deacon, Diana, Sean Hewitt, Chien-Ming Yang, and Masanouri Nagata. 2000. Event-related potential indices of semantic priming using masked and unmasked words: evidence that the n400 does not reflect a post-lexical process. Cognitive Brain Research 9:137–146.

Ettinger, Allyson, Naomi Feldman, Philip Resnik, and Colin Phillips. 2016. Modeling n400 amplitude using vector space models of word representation. In CogSci.

Federmeier, Kara D, and Marta Kutas. 1999. A rose by any other name: Long-term memory structure and sentence processing. Journal of memory and Language 41:469–495.

Firth, John R. 1957. A synopsis of linguistic theory, 1930-1955, Studies in linguistic analysis.

Hagoort, Peter, Lea Hald, Marcel Bastiaansen, and Karl Magnus Petersson. 2004. Integration of word meaning and world knowledge in language comprehension. science 304:438–441.

Harris, Zellig S. 1954. Distributional structure. Word 10:146–162.

Holcomb, Phillip J. 1993. Semantic priming and stimulus degradation: Implications for the role of the n400 in language processing. Psychophysiology 30:47–61.

Huth, Alexander G., Wendy A. de Heer, Thomas L. Griffiths, Frédéric E. Theunissen, and Jack L. Gallant. 2016. Natural speech reveals the semantic maps that tile human cerebral cortex. Nature 532:453.

Khaligh-Razavi, Seyed-Mahdi, and Nikolaus Kriegeskorte. 2014. Deep supervised, but not unsupervised, models may explain IT cortical representation. PLoS computational biology 10:e1003915.

Kiefer, Markus. 2002. The n400 is modulated by unconsciously perceived masked words: Further evidence for an automatic spreading activation account of n400 priming effects. Cognitive Brain Research 13:27–39.

Kriegeskorte, Nikolaus, and Rogier A. Kievit. 2013. Representational geometry: Integrating cognition, computation, and the brain. Trends in cognitive sciences 17:401–412.

Kutas, Marta, and Kara D. Federmeier. 2011. Thirty years and counting: finding meaning in the N400 component of the event-related brain potential (ERP). Annual review of psychology 62:621–647.

Kutas, Marta, and Steven A. Hillyard. 1989. An electrophysiological probe of incidental semantic association. Journal of Cognitive Neuroscience 1:38–49.

Lehmann, Dietrich, and Wolfgang Skrandies. 1980. Reference-free identification of components of checkerboard-evoked multichannel potential fields. Electroencephalography and Clinical Neurophysiology 48:609–621.

Lin, Dekang. 1998. An information-theoretic definition of similarity. In ICML, volume 98, 296–304. Citeseer.

Lorch Jr, Robert F. 1982. Priming and search processes in semantic memory: A test of three models of spreading activation. Journal of verbal learning and verbal behavior 21:468–492.

Mikolov, Tomas, Kai Chen, Greg Corrado, and Jeffrey Dean. 2013. Efficient estimation of word representations in vector space, arxiv preprint arxiv:1301.3781.

Miller, George A. 1986. Dictionaries in the mind. Language and cognitive processes 1:171–185.

Neely, James H. 1976. Semantic priming and retrieval from lexical memory: Evidence for facilitatory and inhibitory processes. Memory & Cognition 4:648–654.

Neely, James H. 1977. Semantic priming and retrieval from lexical memory: Roles of inhibitionless spreading activation and limited-capacity attention. Journal of experimental psychology: general 106:226.

Pennington, Jeffrey, Richard Socher, and Christopher Manning. 2014. GloVe: Global vectors for word representation. In Proceedings of the 2014 conference on empirical methods in natural language processing (EMNLP), 1532–1543.

Poeppel, David. 2012. The maps problem and the mapping problem: two challenges for a cognitive neuroscience of speech and language. Cognitive neuropsychology 29:34–55.

Pylkkänen, Liina. 2019. The neural basis of combinatory syntax and semantics. Science 366:62–66.

Resnik, Philip. 1995. Using information content to evaluate semantic similarity in a taxonomy.

Resnik, Philip. 1999. Semantic similarity in a taxonomy: An information-based measure and its application to problems of ambiguity in natural language. Journal of Artificial Intelligence Research 11:95–130.

Roelofs, Ardi. 1992. A spreading-activation theory of lemma retrieval in speaking. Cognition 42:107–142.

Turney, Peter D., and Patrick Pantel. 2010. From frequency to meaning: Vector space models of semantics. Journal of Artificial Intelligence Research 37:141–188.

Wang, Xuefei, Hao Zhu, and Xing Tian. 2019. Revealing the temporal dynamics in non-invasive electrophysiological recordings with topography-based analyses. biorxiv preprint. https://doi.org/10.1101/779546.

Yamins, Daniel, Ha Hong, Charles Cadieu, Ethan Solomon, Darren Seibert, and James DiCarlo. 2014. Performance-optimized hierarchical models predict neural responses in higher visual cortex. Proceedings of the National Academy of Sciences 111:8619–8624.

Yang, Jinbiao, Hao Zhu, and Xing Tian. 2018. Group-level multivariate analysis in EasyEEG toolbox: Examining the temporal dynamics using topographic responses. Frontiers in Neuroscience 12:468.

Zhang, Linmin, and Liina Pylkkänen. 2015. The interplay of composition and concept specificity in the left anterior temporal lobe: An meg study. NeuroImage 111:228–240.

